# It’s a wrap: deriving distinct discoveries with FDR control after a GWAS pipeline

**DOI:** 10.1101/2025.06.05.658138

**Authors:** Benjamin B. Chu, Zihuai He, Chiara Sabatti

## Abstract

The standard analysis pipeline for genome-wide association studies (GWAS) is based on marginal tests of association. These are computationally convenient and portable, but the discoveries resulting from their rejections are not immediately interpretable, and require post-processing as “clumping” and “fine mapping.” An interesting alternative is provided by conditional independence hypotheses: their rejections lead to the identification of distinct signals across the genome, accounting for measured confounders, and pointing to separate causal pathways.

An obstacle to the wide adoption of this approach has been that it requires access to individual level data. Overcoming this barrier, recent work has shown how summary statistics resulting from the standard marginal GWAS analysis can be used as input of a procedure to test conditional independence hypotheses while controlling the false discovery rate. This secondary analysis requires sampling of synthetic negative controls (knockoffs) from a distribution determined by the linkage disequilibrium patterns in the genome of the population under study. In prior work, we have pre-computed this distribution for European genomes, starting from information derived from the UK Biobank. Thus, researchers working with GWAS in a European population can carry out a knockoff analysis with minimal computational costs, using the distributed routine `GhostKnockoffGWAS`.

Here we introduce and release a new software (`solveblock`) that extends this capability to a much richer collection of studies. Given a set of genotyped samples, or a reference dataset, our pipeline efficiently estimates the high-dimensional correlation matrices that describe dependencies across the genome, making rather common sparsity assumptions. Taking this sample-specific estimate as input, the software identifies groups of genetic variants that are highly correlated, and uses them to define an appropriate resolution for conditional independence hypotheses. Finally, we compute the distribution for the exchangeable negative controls necessary to test these hypotheses. The output of `solveblock` can be passed directly to `GhostKnockoffGWAS`, allowing users to carry out the complete analysis in a two step procedure.

Simulations, based on five UK Biobank sub-populations, illustrate the method’s FDR control. The analysis of 26 phenotypes of varying polygenicity in British individuals, results in `≈` 19 additional discoveries, compared to standard marginal association testing. Our code, precompiled software, and processed files for these five sub-populations are openly shared.

## 1 Introduction

The knockoff filter [1] is a variable selection framework that tests conditional independence hypotheses while controlling the false discovery rate (FDR) [2]. The approach is particularly appealing in genome-wide association studies (GWAS), where it leads to interpretable discoveries, that do not require post-processing and are closely tracking causal variants [3]. A successful implementation of this strategy models the distribution of genotypes using hidden-Markov models (HMM) [4, 5], in line with the genetic literature. Prior to a knock-off analysis, however, this approach requires phasing the genotypes, estimating parameters of an HMM, and determining identical-by-descent segments. All these steps are computationally intensive, despite recent advances [6, 7, 8], and require reliance on a collection of external tools not optimized for this specific goal. As a result, a knockoff analysis of GWAS data based on HMMs is still much more involved than the traditional GWAS pipeline.

An alternative path to testing conditional hypotheses across the genome has started to emerge with the realization that an approximate knockoffs analysis can be carried out working not with individual level data, but with summary statistics capturing the correlation between the outcome and the explanatory variables [9, 10]. This procedure, known as GhostKnockoffs, takes as input GWAS summary statistics (e.g. p-value and direction of effect) and can therefore be run without changing any of the standard pre-processing steps. Empirical studies [11] show a particularly appealing characteristic of this pipeline: the input can be any testing statistics that is appropriate for the data, such as those derived from linear mixed models that account for relatedness, as long as its null distribution is *N*(0, 1). In [12] the re-analyses of 60+ European GWASes with this framework lead to greater power and improved precision, as compared with the results of the original analyses.

While the GhostKnockoffs approach has the potential of a more straightforward harmonization with the standard GWAS pipeline, the necessity of modeling the distribution of genotypes and calculating the derived knockoff distribution remains. Since this approach relies on matching of the first two moments, crucial elements are (a) the matrix of correlations between allelic counts of different SNPs and (b) the sampling of powerful second order knockoffs [13]. The authors of [12] distributed a software (`GhostKnockoffGWAS`) that takes as input GWAS summary statistics and parameters that characterize the knockoff sampling distribution (a,b). To facilitate users, pre-computed parameters appropriate for a European populations derived from LD matrices calculated on data from the UK Biobank [14, 15], were also made available on Zenodo [16].

In this work we extend the access to a computationally convenient and reliable knockoff analysis to a much richer set of studies. Given that GhostKnockoff can be used as a “wrapper” around any standard pipeline, what remains is to provide users with a way of computing the parameters that characterize the knockoff sampling distribution (a,b) that are appropriate for their specific sample. The software we introduce assumes that individual-level genotype data is available. This can be from the sample understudy or an appropriate reference set. Taking this as input, it (1) estimates high-dimensional LD matrices appropriate for the target sample; (2) it identifies groups of SNPs in high disequilibrium that will be the object of conditional inference; (3) it approximates the dependency structure across these groups; and (4) computes the parameters of the joint variance covariance matrix of the original variables and knockoffs to maximize power. Note that if an adequate LD matrix is available (instead of individual level data), step (1) can be skipped; one of the purposes of the release software is to facilitate (1) when this is not the case.

Thanks to the `solveblock` binary executable, users can construct the bundle of LD files necessary for GhostKnockoff analysis directly from individual level data stored in VCF [17] or binary PLINK [18] format. To run a genomewide knockoff-based conditional independence testing, then, users need to (i) run any standard GWAS pipeline to obtain marginal Z-scores, (ii) run the `solveblock` executable to obtain parameters for knockoff construction, and (iii) input the results of these two first steps into `GhostKnockoffGWAS`. Users have the option (iv) to share the output of `solveblock` by uploading it to services such as Zenodo [16]. Simulation studies and real-data examples illustrate the speed, enhanced power, and FDR control of knockoff-based inference enabled by this pipeline.

## 2 Methods

We assume that the user has available a set of genotypes **X** on *n* individuals (rows) and *p* SNPs (columns) as well as summary statistics on the association between these same SNPs and a phenotype of interest. The genotypes might have been obtained as part of the study that lead to the association summary statistics, or belong to an appropriate reference panel. We note that when individual level data on genotype and phenotype is available, one can conduct a knockoff-based analysis by generating knockoffs 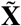 using HMMs distribution [4, 3, 5]. While there are many robustness advantages of this approach, it remains more computationally involved, and it is more difficult to harmonize with analysis pipelines that might have been pre-established, as it is often the case in the context of consortia. In what follows, we work with a GhostKnockoff [10] pipeline [12].

Individual-level data (VCF or binary PLINK format)

### Preprocessing steps

For GWAS data with *p* `≈` 10^6^, it is common to assume that the variance-covariance matrix **Σ** for the SNPs has a block-diagonal form [19, 20]:

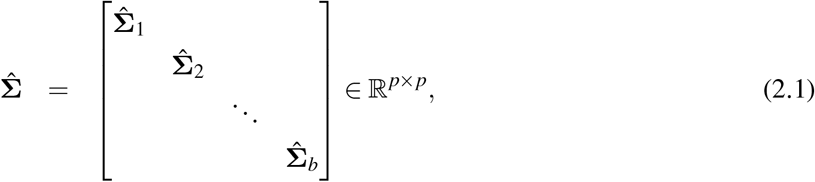

effectively assuming that SNPs in different blocks are independent. Defining blocks in correspondence to different chromosomes is certainly natural. However, in order to reduce computational burden, researchers typically consider blocks of smaller sizes. It is important to recall that if the block-wise independence reflected in (2.1) is not (at least approximately) satisfied, the knockoff analysis resulting from the pipeline we are about to describe will not have FDR control.

The block boundaries can be computed efficiently using the `snp_ldsplit` function of the `bigsnpr` package [20] and need to be identified before `solveblock` can be run. Empirically, working in a variety of populations, we found that *b* on the order of 10^2^ ∼ 10^3^ is sufficient to partition the human genome based on the default objective provided in `bigsnpr`, see Table 1. To obtain reliable estimates to genetic correlation, we recommend that only SNPs with minor allele frequency `≥` 0.01 should contribute to LD block estimation. Note that, while more numerous than the chromosomes, these blocks include a large number of SNPs, so that they can be used to represent a fairly extensive linkage disequilibrium.

**Table 1:** Summary of various LD files after applying the snp_ldsplit function of bigsnpr package to the individual-level data of UK-Biobank.

| Data | Sample size ($n$) | Number of blocks ($b$) | disk space (GB) |
| --- | --- | --- | --- |
| British | 306604 | 636 | 20.0 |
| Indian | 5951 | 615 | 13.0 |
| Caribbean | 4517 | 489 | 13.0 |
| African | 3394 | 513 | 9.4 |
| Chinese | 1574 | 505 | 7.4 |

Within each block *i* = 1, …, *b* in equation (2.1), `solveblock` will perform 4 main steps, summarized in Figure 1, and described in the following.

**Figure 1:**
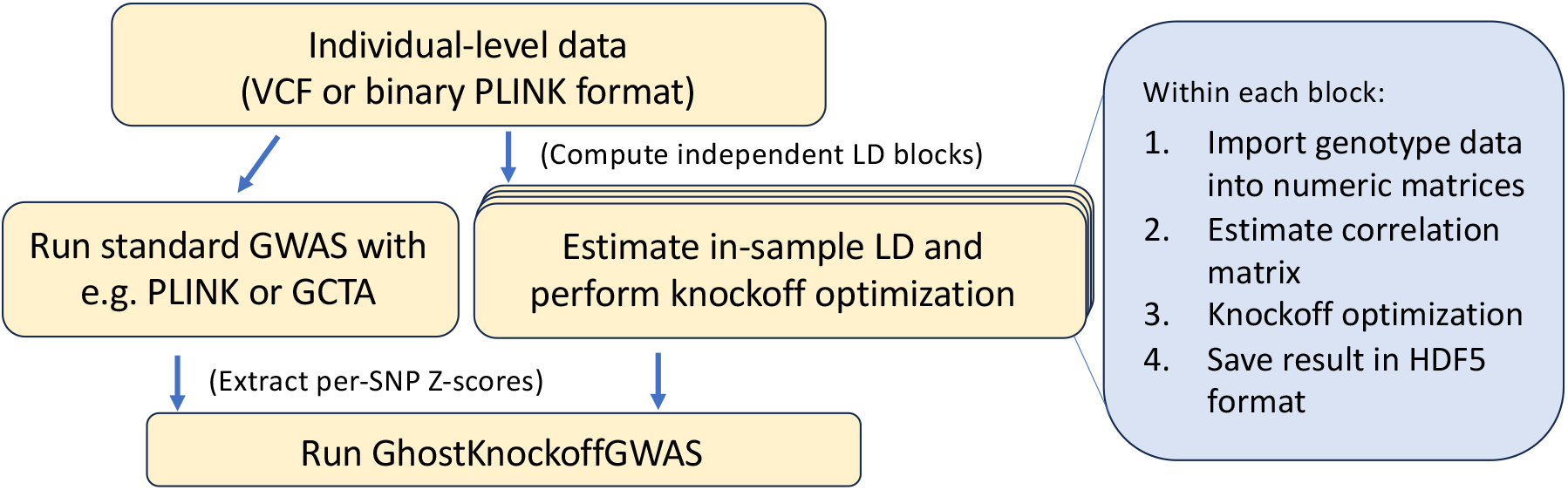
Overview of the GhostKnockoff pipeline. The sequence of events taking place within the `solevblock` executable are outlined in the blue box.

### 2.1 Import genotype data into computer memory

We import the genotype data into double-precision matrices *x*_*ij*_ ∈ {0, 1, 2, missing} using the `OpenMendel` [21] modules `SnpArrays.jl` or `VCFTools.jl`. This step requires 8*np*_*i*_ bytes of computer RAM, which is memory intensive if *p*_*i*_ (number of variants in block *i*) is extremely large. Typically, however, the block-diagonal approximation (2.1) implies *p*_*i*_ is generally on the order of 10^3^, which can be handled efficiently.

For VCF inputs, our convention is to convert each `ALT` allele into numeric value of 1. With binary PLINK inputs, all `A2` alleles are converted into 1 even if it is the major allele (note this is equivalent to using `–-keep-allele-order` within `PLINK 1.9` software). Multi-allelic SNPs should be split prior to analysis. Missing genotypes are automatically imputed by column mean.

### 2.2 Estimation of correlation matrix

Within region *i* ∈ {1,…, *b*}, let 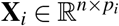 denote the column-standardized genotype matrix. Our next goal is to estimate a correlation matrix. Note that the pre-processing step of identifying block boundaries technically is based on measures of correlation, implicitly used by `snp_ldsplit` function of `bigsnpr` package. Once the boundaries are identified, however, we explicitly estimate a correlation matrix (denoted by 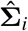) that will be used in the GhostKnockoff pipeline. We adopted the same LD estimation routine of Pan-UKB [15] which we found to perform well [13]. Letting **C** denote a matrix of covariates (sex, age, etc), we compute:

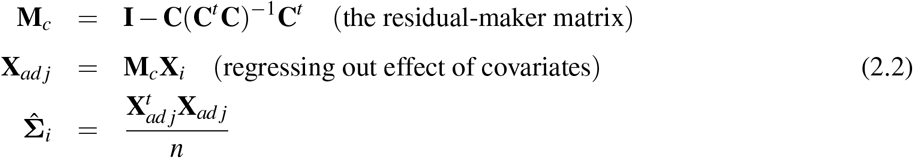

For numerical stability, we set the minimum eigenvalue of 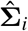 to be 0.00001 and scaled the final result so the diagonal entries are exactly 1. (Inputting the the matrix of covariates **C** is optional).

### 2.3 Parameters for group-knockoffs generation

Our next step is to calculate the parameters that describe the variance-covariance matrix of the original variables and knockoffs. For each block **Σ**_*i*_, this includes 3 main steps: (1) identify groups of highly correlated SNPs that are going to be the basis of inference, (2) approximate the joint distribution of all SNPs using a conditional independence model in which only a subset of “key variables” within each group are driver of the correlations across groups, and (3) find the variance-covariance matrix for 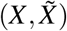 that satisfies the exchange-ability constrains while maximizing detection power by maximizing the entropy criteria [22]. The motivation behind all of these steps and the algorithms required to carry them out are described elsewhere [13]. We summarize them below for completeness.

Following [13], for each 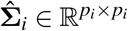, we define non-overlapping groups via average-linkage hierarchical clustering with correlation cutoff 0.5. Given group boundaries, we then apply the best subset selection algorithm of [13] (Algorithm A2 in the supplement) with *c* = 0.5 to identify *group-key* variables within each group. Intuitively, the group-key variables are a subset of {1,…, *p*_*i*_} that drive the dependence across groups. The sampling of knockoffs, then, can be carried out by sampling knockoffs for these key variables and then, within each groups, knockoffs for the remaining variables conditional of these. Let *R* denote the set of group-key variables, *C* denote its complement, and 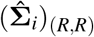 the rows and columns of 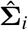 corresponding to variables in *R*. Knockoff optimization proceeds by solving the maximum entropy [22] objective

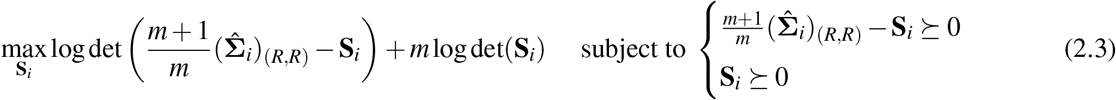

where *m* = 5 is the number of knockoffs to generate per feature and **S**_*i*_ ⪰ 0 means **S**_*i*_ is positive semi-definite. Then the *p*_*i*_ × *p*_*i*_ matrix

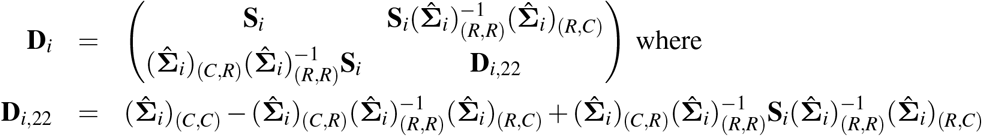

is the solution to the maximum entropy objective for the full 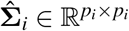. In particular, **D**_*i*_ ⪰ 0 satisfies the constraints 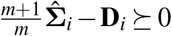 and can be used for ghost-knockoff sampling [23]. We used the coordinate descent algorithm implemented in `Knockoffs.jl` [13] to solve equation (2.3). Note that this procedure generates second order knockoffs, which coincide with full fledged knockoffs when the genotypes 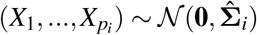 [24]. While this latter assumption is unrealistic, second-order approximations lead to effective FDR control when second order statistics are at the center of the data analysis methods—this is the case for standard GWAS pipelines.

### 2.4 Output in HDF5 format

Up until this point, we have assembled the reduced data:

- 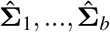: correlation matrices for each block
- **S**_1_, …, **S**_*b*_: knockoff optimization matrices for each block, carried out on the group-key variables
- **D**_1_, …, **D**_*b*_: knockoff sampling matrices for each block satisfying 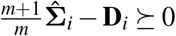.
- Group membership vector for each block *i* = {1,…, *b*}
- Group-key variables for each block *i* = {1,…, *b*}

We will save this ensemble of data in HDF5 format [25] which can be directly read by `GhostKnockoffGWAS` [12]. Note that due to the block-diagonal partitioning in equation (2.1), this bundle of “pre-processed” LD files are typically dozens of gigabytes in size, and contain no individual level data. We emphatise this last point to invite researchers to share their results as they may be used by other studies on similar populations.

## 3 Result

To illustrate the software, we are going to leverage simulations and real data analysis. Both are based on data obtained from the UK-Biobank. We start by describing the data we use and its pre-processing steps.

### 3.1 UK-Biobank (UKB) data

#### Quality Control

The second release of the UK Biobank (UKB) data [14] contains ∼ 500, 000 samples of primarily European descent and ∼ 800, 000 SNPs without imputation. Following [3], for British samples, we kept only 1 sample from each of the 60,169 familial groups. This resulted in *n* = 306, 604 unrelated British samples. Next, we also created different sub-populations based on self-reported Indian (*n* = 5951), Caribbean (*n* = 4517), African (*n* = 3394), and Chinese (*n* = 1574) ancestry. For these non-British sub-populations, we kept related samples to mimic cryptic relatedness in downstream simulations. For all five UKB sub-populations, we excluded non-biallelic SNPs with minor allele frequency (MAF) < 0.01 and Hardy-Weinburg equilibrium p-values < 10^−6^. Finally, for our real-data analysis, we chose 26 continuous and commonly measured phenotypes (see Table 2) as targets of analyses. Samples that cannot be matched to phenotype file by ID or whose phenotypes contain missing values were also excluded.

**Table 2:**
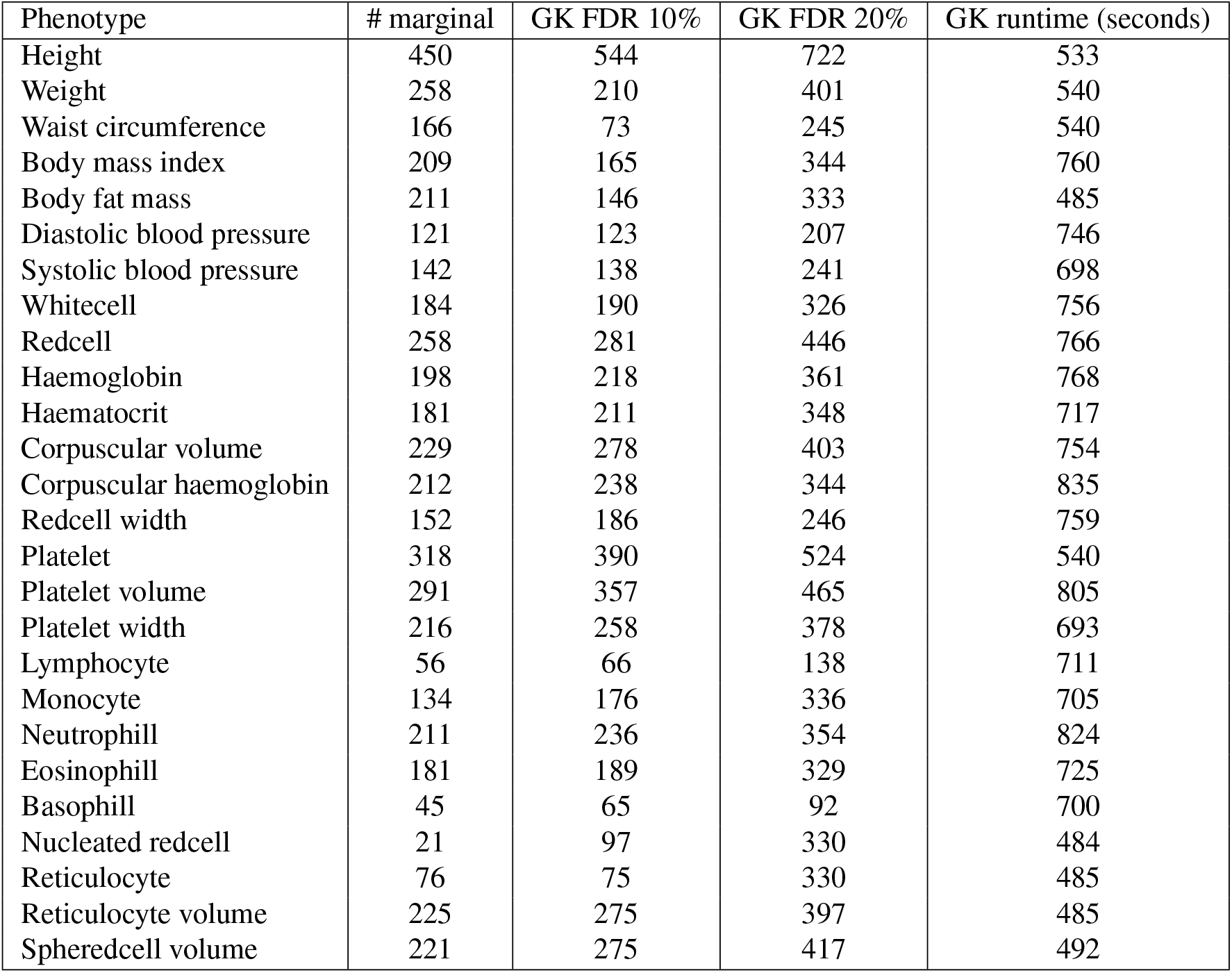
The number of independent discoveries for various phenotypes analyzed under both the knockoff framework and the conventional marginal hypothesis testing. GK stands for ghost-knockoff and FDR stands for false-discovery rate. The target FDR is set to 10% and 20%. Note the runtime corresponds to the runtime of `GhostKnockoffGWAS` and does not include the runtime for `solveblock`, since the later only needs to be done once for a population of interest.

#### Non-genetic covariates

Following Pan-UKB [15], we included the following covariates for LD adjustments: sex, age, age^2^, sex×age, sex×age^2^, and the top 10 principal components precomputed by the UK-Biobank. These covariates form the matrix **C** in equation (2.2).

### 3.2 LD file Construction

Our first goal is to construct the necessary LD files for various subpopulations of the UKB, to enable knockoff-based analysis and simulations on these populations. To do so, we first applied the `snp_ldsplit` functionality of `bigsnpr` package [20] to each chromosome of each subpopulation of UKB separately. We used the parameters `thr_r2` = 0.01, `max_r2` = 0.3, `min_size` = 500, and `max_size` = {1000, 1500, 3000, 6000, 10000}. Given block boundaries in each subpopulation, we applied our `solveblock` executable directly on the individual-level genotypes. We submitted a separate cluster job for each region, and every job finished within a few minutes. Table 1 summarizes the final number of blocks and disk space needed to store the resulting LD files.

### 3.3 Simulations: FDR control

In this section, we demonstrate the proposed pipeline results in valid (group) FDR control when the output of `solveblock` is directly used by `GhostKnockoffGWAS` [12]. We restrict our attention to SNPs falling on chromosome 22, totaling *p* = 8878 SNPs. In all simulations we column-standardize **X** and simulate the response as **y** = **X*β*** + `N`(**0**, 3**I**_*n*×*n*_) where ***β*** contain *k* = 10 non-zero effects chosen randomly and effect sizes drawn from `𝒩` (0, 0.5). The narrow sense heritability for these simulations is approximately 0.25. After regressing out the effect of covariates, we standardize the phenotype.

Marginal Z-scores were computed in two ways. For the British samples, where related individuals are removed, we compute marginal Z-scores by the score statistic 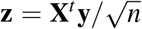 where **y** is standardized. For other populations in the UKB, which contain related samples and potential continental admixture, we use the linear mixed model software GCTA [26] to compute p-values, and transformed them back into Z-scores. Once LD files are estimated from genotypes, and Z-scores evaluated, they are used as the only inputs used for `GhostKnockoffGWAS` for evaluating power and (group) FDR.

The results of 200 independent simulations are depicted in Figure 2. The procedure consisting in `solveblock` + `GhostKnockoffGWAS` results in (group) FDR control. The variation of power across all populations is a consequence of variable sample size, as phenotypes are generated from the same distribution.

**Figure 2:**
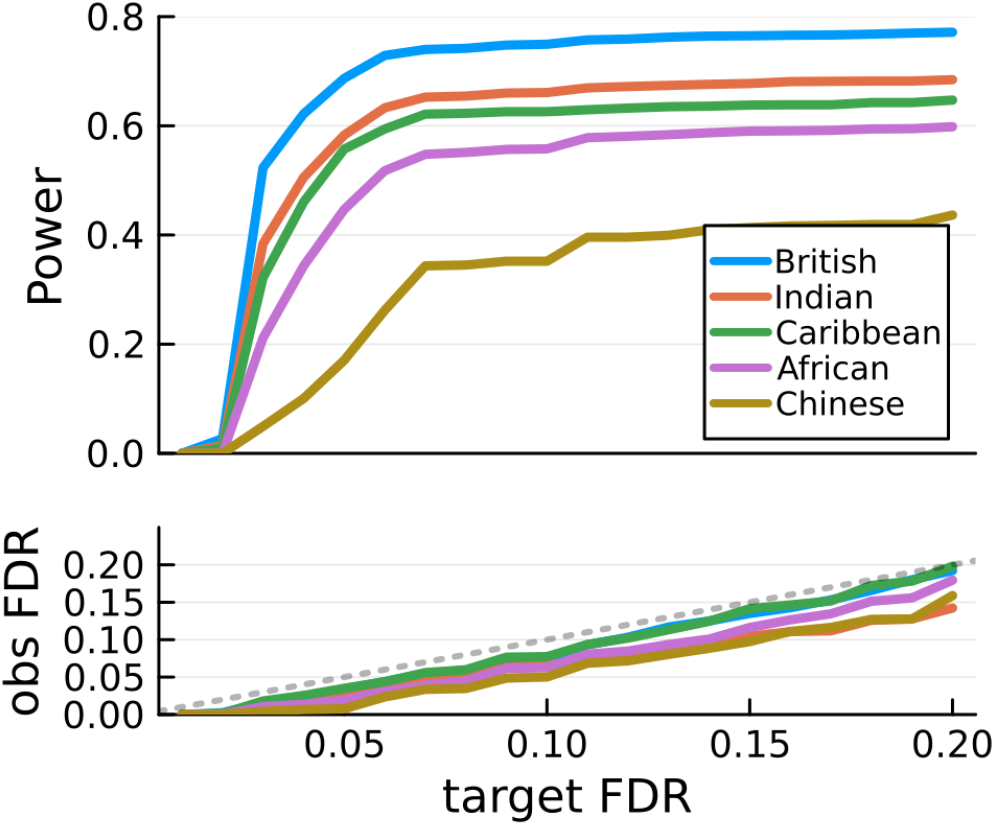
Average power and FDR across 200 simulations. The British subset contains only unrelated individuals. For other populations, related samples are included in the analysis.

### 3.4 Real data analysis: 26 UKB phenotypes using GhostKnockoff

Here we analyzed 26 continuous phenotypes based on the British samples of UKB. The goal of this section is to (1) demonstrate the scalability of GhostKnockoff on biobank-scale datasets (each phenotype took <15 minutes to run on a single CPU), and (2) demonstrate the enhanced power of GhostKnockoff in locus discovery compared to marginal analysis methods.

Because `GhostKnockoffGWAS` relies on marginal Z-scores as inputs, we first conduct a standard marginal analysis using the score statistic after regressing out the effect of covariates from the phenotype vector. Mathematically, let **y** be the original phenotype vector, **C** be the matrix of covariates, and **X** the genotype matrix that is column standardized to mean 0 variance 1. We first compute 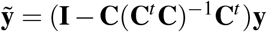 and standardize it to mean 0 variance 1. Then we compute 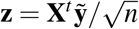 where *n* = 306, 629 British samples. The resulting **z** in combination with the output of `solveblock` are then used as inputs for `GhostKnockoffGWAS` [12].

Figure 3 compares the number of independent discoveries from knockoff-based analyses to that from conventional marginal GWAS with Bonferroni threshold set to 5 × 10^−8^. While each knockoff discovery is by construction distinct, for ease of comparison, 2 discoveries (coming from marginal or knockoff analyses) are considered *independent* if they are at least 1 Mb apart. Based on this criteria, ghost knockoff analyses found 19.0 (149.1) additional independent discoveries compared to marginal testing when the target FDR is 10% (20%). Table 2 provides more details. Notably, ghost-knockoff discovered more signals in 20 out of 26 traits given target FDR of 10%, and each phenotype took 15 minutes or less to run on a single CPU core on the cloud. These result showcases our method’s enhanced power and computational efficiency for locus discovery.

**Figure 3:**
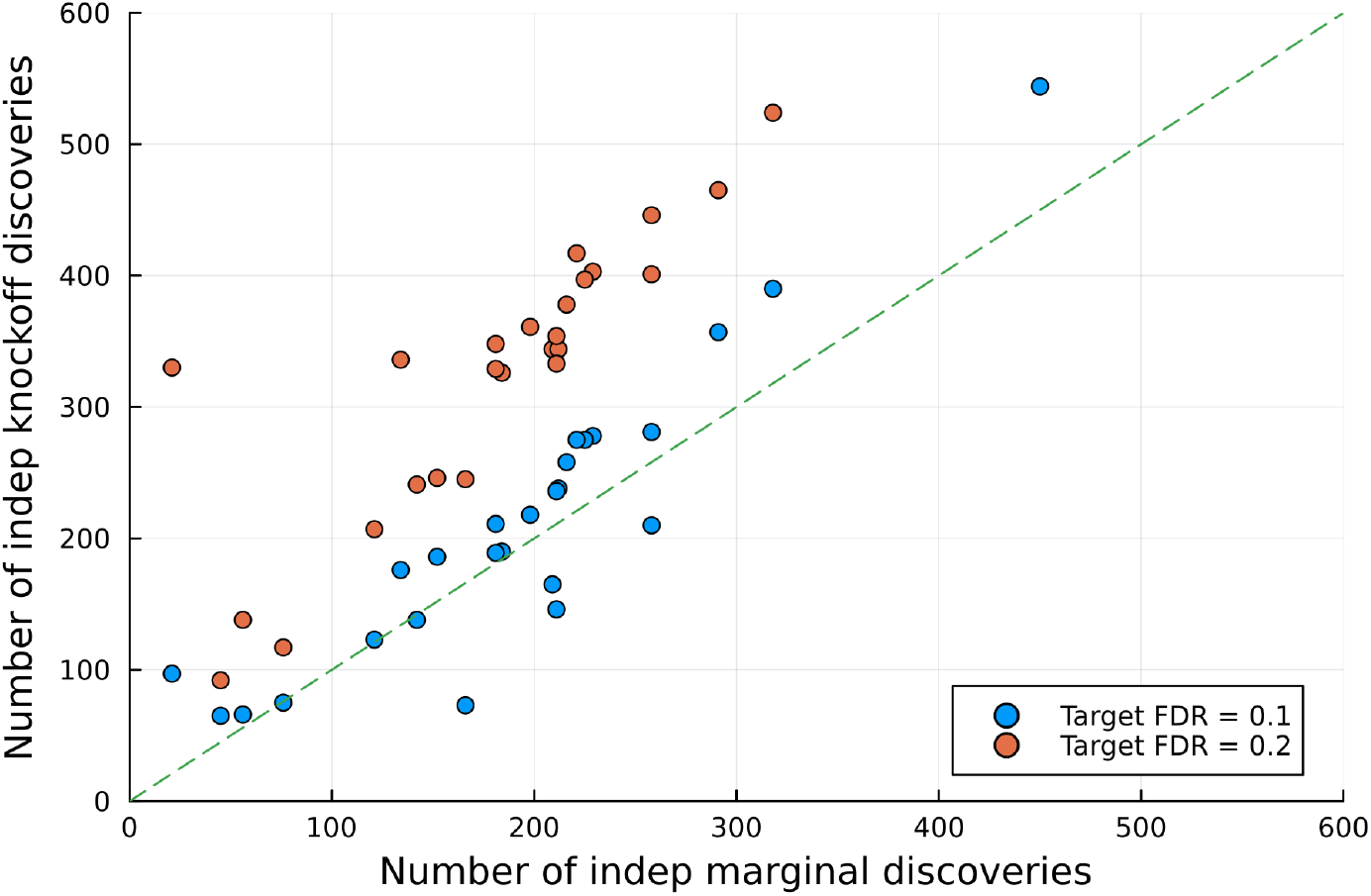
The number of (independent) discoveries from GhostKnockoff compared to a conventional marginal GWAS. Each dot is a different phenotype (26 total), each of which took <15 minutes on a single CPU to run. Dots above the dashed line indicate that knockoff analysis discovered more signals compared to marginal analysis. Details in Table 2.

## 4 Discussion

The goal of this paper is two fold. First, we introduce the `solveblock` executable. When combined with the accompanying software `GhostKnockoffGWAS` [12], this can be used to perform an end-to-end GWAS analysis using a knockoff perspective while adhering to a standard GWAS pipeline. We found that this approach is scalable to biobank-sized data and controls the FDR, while delivering enhanced power.

The second goal of this paper is to provide a platform for and encourage data exchange. Prior to this work, precomputed LD parameters necessary for GhostKnockoff analyses were available only for European samples. We now provide these for Indian, Caribbean, African, and Chinese samples, with the caveat that the parameters have been constructed from the fairly small sample sizes available in the UK biobank. More impactfully, the `solveblock` executable introduced in this paper will allow investigators to generate parameters that reflect the LD in their specific samples. By construction, the output files are small in size (Table 1), do not contain private information, and are therefore easy to share. For example, the results can be easily uploaded to Zenodo [16]. If just a small fraction of our users are open to data-sharing, `GhostKnockoffGWAS` will become exponentially more versatile.

Further extensions are also planned. First, we note that all ghost-knockoff computation can feasibly be carried out entirely on the cloud. We then envision the construction of a utility similar to the Michigan’s imputation server [27]: users would upload GWAS summary statistics, select the desired LD panel to use, and later download the ghost-knockoff discoveries. This bypasses the need to distribute processed LD files and avoids headaches with software installation. Next, it is now realistic to apply the `solveblock` executable to other datasets, such as the All of US [28] and Million Veteran Project [29]. Enabling GhostKnockoff analyses on these newer biobanks will potentially increase the power of thousands of related projects, but also improve health disparities for minorities who are not well represented in the UK Biobank.

Finally, it is of interest to extend the ghost-knockoff methodology to non-SNP-array data, such as those generated from high-coverage whole genome sequencing or low-pass imputation-based data [30]. Such dense genotyping technologies are expected to become increasingly affordable, eventually leading to a higher adoption rate than array-based data in genetic studies. While the simulation studies here show empirical FDR control, the current GhostKnockoff pipeline is likely too slow for analyzing high-coverage sequenced data, since the blocks in equation (2.1) routinely have dimensions exceeding 10^5^. In such cases, more computational innovation is needed for knockoff optimization (2.3) and GhostKnockoff sampling [12]. We raise these research questions to stimulate discussions and encourage participation from the broader community.

## 5 Web Resources

**Project home page**: https://github.com/biona001/GhostKnockoffGWAS

**Supported operating systems**: Linux

**Programming language**: Written in Julia [31] but program is compiled into standalone executables

**License**: MIT

**Reproducibility scripts**: The code used to run the simulated experiements and real data analyses can be found in https://github.com/biona001/solveblock-reproducibility.

## 6 Acknowledgments

B.C. and C.S. were supported by the grants R01MH113078, R56HG010812, NSF 2210392, and R01MH123157. B.C. was additionally supported by the NLM T15 Postdoc training grant. Z.H. were supported by NIH/NIA awards AG066206 and AG066515. Access to UK Biobank data was through Application 27837.

